# Spatial Transcriptomics Prediction from Histology jointly through Transformer and Graph Neural Networks

**DOI:** 10.1101/2022.04.25.489397

**Authors:** Yuansong Zeng, Zhuoyi Wei, Weijiang Yu, Rui Yin, Bingling Li, Zhonghui Tang, Yutong Lu, Yuedong Yang

## Abstract

The rapid development of spatial transcriptomics allows for the measurement of RNA abundance at a high spatial resolution, making it possible to simultaneously profile gene expression, spatial locations, and the corresponding hematoxylin and eosin-stained histology images. Since histology images are relatively easy and cheap to obtain, it is promising to leverage histology images for predicting gene expression. Though several methods have been devised to predict gene expression using histology images, they don’t simultaneously include the 2D vision features and the spatial dependency, limiting their performances. Here, we have developed Hist2ST, a deep learning-based model using histology images to predict RNA-seq expression. At each sequenced spot, the corresponding histology image is cropped into an image patch, from which 2D vision features are learned through convolutional operations. Meanwhile, the spatial relations with the whole image and neighbored patches are captured through Transformer and graph neural network modules, respectively. These learned features are then used to predict the gene expression by following the zero-inflated negative binomial (ZINB) distribution. To alleviate the impact by the small spatial transcriptomics data, a self-distillation mechanism is employed for efficient learning of the model. Hist2ST was tested on the HER2-positive breast cancer and the cutaneous squamous cell carcinoma datasets, and shown to outperform existing methods in terms of both gene expression prediction and following spatial region identification. Further pathway analyses indicated that our model could reserve biological information. Thus, Hist2ST enables generating spatial transcriptomics data from histology images for elucidating molecular signatures of tissues.

## 1. Introduction

Spatially resolved transcriptomics (ST) technologies have recently bloomed as the extended technologies of single-cell RNA sequencing (scRNA-seq), such as Slide-seq [1], XYZeq [2], and 10X Visium. STs hold the potential of profiling gene expression for the whole transcriptomics at nearly single cell resolution accompanied with their spatial locations, for many techniques, and often with the matched whole slide of hematoxylin and eosin (H&E)-stained histology images (WSI) [3–5]. The sequenced expressions at each spot contain two to dozens of cells depending on different ST technologies [6]. These powerful ST techniques have transformed our views on the developing human heart, amyotrophic lateral sclerosis, and Alzheimer’s disease [7–9]. However, it remains a challenge for current available computational methods by incorporating the unique properties of spatial transcriptomics data to unveil spatially variable features and cell identities [10]. To take full advantage of the added spatial information, several novel computational methods have been developed for exploring the spatial expression patterns (SPARK [11], SpatialDE [12], and CoSTA[13], etc.), clustering spatial domains (stLearn [14], SpaGCN [15], BayesSpace [16], and STAGATE [17], etc.), analyzing the cell type compose of spots (DSTG [6], SPOTlight [18], Giotto[19], and SpatialD-WLS [20], etc.), learning the spatial embedded representation (SEDR [21], conST [22], and MAPLE[23], etc.), and inferring cell-cell communication (SpaOTsc [24], DistMap [25], and Tangram[26], etc.).

Despite several computational methods that have been designed for the enriched information brought by ST technologies, ST technologies haven’t been utilized in large-scale studies due to the expensive costs. Relatively, WSIs are much cheaper and easier to obtain and are routinely generated in clinics [27]. In fact, WSIs have been shown highly correlated with gene expression, and have been used to predict bulk gene expression profiles through HE2RNA[28]. HE2RNA could robustly capture subtle structures in WSIs, unraveling an enlightening tumor region for the development of specific cancer types. Similarly, WSIs have also been used to predict spatial gene expressions. For example, ST-Net [29] predicts spatially variable gene expression of each spot in the whole histology image, where the histology image patch at the spot is encoded by the DenseNet [30]. The spatially resolved transcriptome predicted by ST-Net enables image-based screening for biomarkers with spatial variation. However, the DenseNet in ST-Net processes the patches independently, their predicted gene expressions are limited to local image patches and can’t obtain global WSI information.

To solve this issue, HisToGene [27] is designed to include spot spatial relations through the Vision Transformer (VIT) [31], where VIT uses a self-attention mechanism to capture the relationships between spots[32]. The attention mechanism has shown stable performance on many tasks such as registration[33] and segmentation [34]. HisToGene outperforms ST-Net in terms of gene expression prediction and clustering tissue regions due to the inclusion of the spot spatial relations. In spite of the inclusion of spatial locations, HisToGene doesn’t capture 2D vision features within the image patch because VIT needs to reshape the patch features into a one-dimensional vector. In addition, the attention mechanism doesn’t explicitly use the neighbored patches that usually play more important roles. Thus, it’s necessary to develop one framework to integrate 2D vision features within the image patch, global spatial relations of patches, and the explicit neighborhood relationships. In fact, the explicit neighborhood relationships can be elegantly solved by graph neural networks (GNN) [35, 36]. GNNs recently attracted much attention due to their ability to efficiently capture structural information. GNNs have been successfully used in many areas, such as biological medicine [37–39] and traffic prediction [40].

In this study, we have developed Hist2ST, a spatial information-guided deep learning method for spatial transcriptomic prediction from WSIs. As shown in Fig 1, Hist2ST consists of three modules: the Convmixer, Transformer, and graph neural network. Hist2ST simultaneously takes account of the 2D vision feature within the image patch and the spatial dependencies among image patches. Specifically, at each sequenced spot, the corresponding histology image is cropped into an image patch. The image patches are fed into the Convmixer module to capture the 2D vision features within the image patch through convolution operations. The learned features are fed into the Transformer module to capture the global spatial dependencies through the self-attention mechanism. Hist2ST then explicitly captures the neighborhood relationships through the graph neural network. Finally, these learned features are used to predict the gene expression by following the zero-inflated negative binomial (ZINB) distribution [41]. To alleviate the impact by the small spatial transcriptomics data, a self-distillation mechanism is employed for efficient learning of the model. Hist2ST was tested on the spatially resolved transcriptomics datasets, and shown to outperform existing methods in terms of both gene expression prediction and spatial regions identification using the predicted expression. Thus, Hist2ST enables generating spatial transcriptomics data from histology images for elucidating molecular signa-tures of tissues.

**Fig 1.**
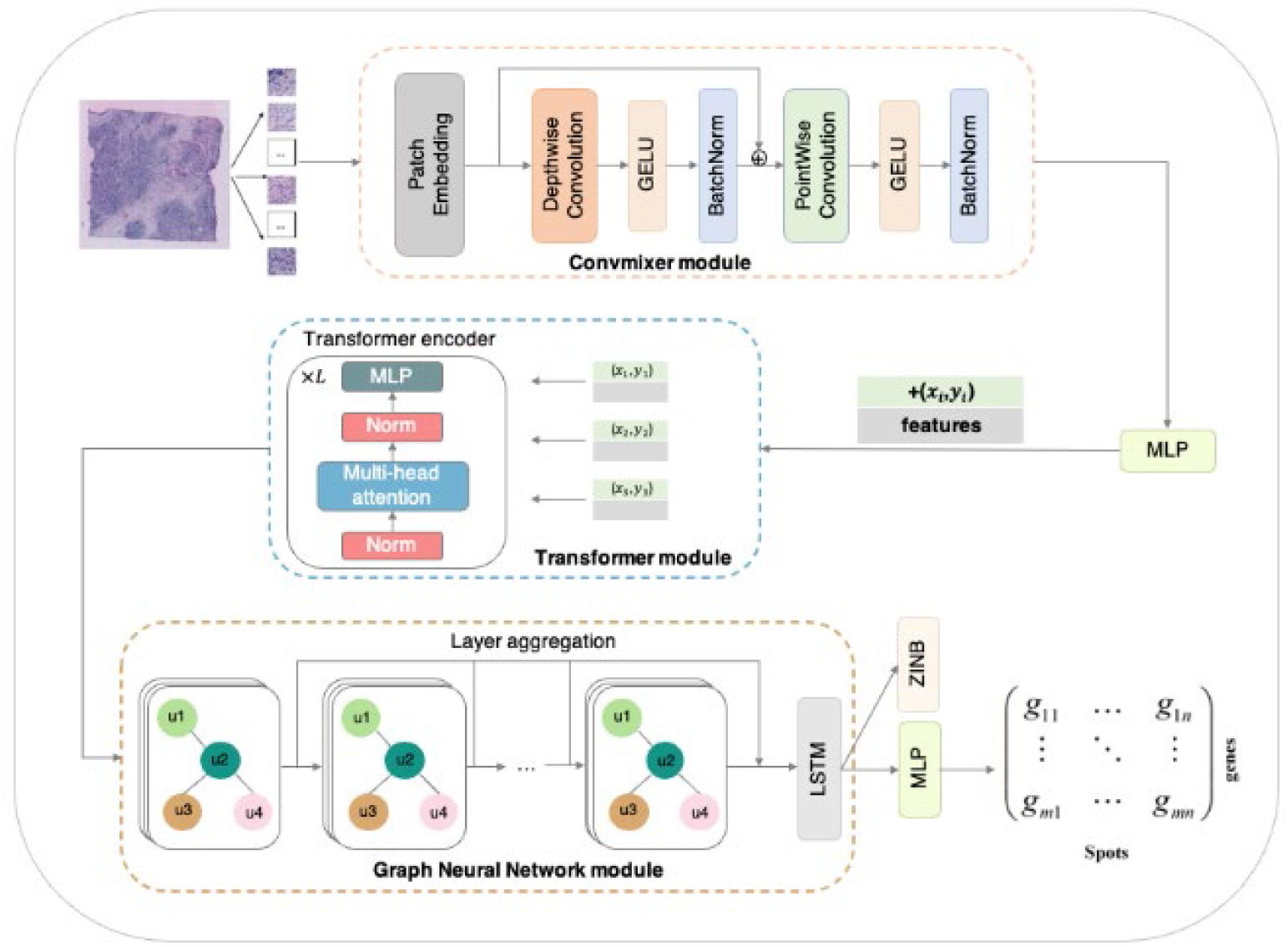
The schematic overview of the Hist2ST for predicting gene expression from histology image. Hist2ST consists of Convmixer, Transformer, and graph neural network modules. Around the sequenced spots, the input histology image is divided into image patches, which are fed into the Convmixer module to capture the 2D vision features. The learned features are then input into the Transformer module to capture the global spatial dependency on the whole image. The outputs are further processed through the graph neural network to capture the local spatial dependency with the neighbor patches. Finally, the learned features are modelled by ZINB distribution to predict the spatial gene expression.

## 2. Materials and Methods

### 2.1 Datasets and pre-processing

We employed the same spatial transcriptomics datasets as the previous study [27]. For completeness, we briefly introduce the procedure to process these datasets, which include the human HER2-positive breast tumor dataset (HER2+) [42] and the human cutaneous squamous cell carcinoma (cSCC) dataset [43]. The HER2+and cSCC datasets consist of histology images, gene expressions at the spatial spots, and their coordinates. The gene expression *X* ∈ *R^N×d^*, where *N* is the number of spots in the histology and *d* is the number of genes. Specifically, the HER2+ breast tumor dataset was measured by the 10x Visium platform including 36 tissue sections from eight patients. We retained 32 sections from seven patients that had at least 180 spots per section. The cSCC dataset contained 12 tissue sections obtained from four patients, with each patient having three sections. The 12 tissue sections of the cSCC dataset were measured by the 10x Visium platform. In the training step of our model, the histology images and the spatial coordinates of the spots will be used as the input, and the matching gene expression data will be used as labels.

We pre-processed the ST datasets. For the histology image, we divided the whole histology image into N × (3 × W × H) patches according to the size (W × H) and location of each spot, where the 3 was the number of channels, and W and H indicated the width and height of the patch, respectively. For all datasets, W and H were set to 112 pixels, which corresponded to the diameter of each spot in the ST data. For the gene expression data of each tissue section, we selected the top 1000 highly variable genes and excluded genes that were expressed in less than 1000 spots across all tissue sections. The feature counts for each spot were divided by the total counts for that spot and multiplied by the 1,000,000. This was then natural-log transformed using log1p.

After pre-processing the ST datasets, HER2+ retained 9,612 spots and 785 genes, and cSCC retained 6630 spots and 134 genes. For evaluating the gene expression prediction accuracy, we conducted leave-one-out cross validation. Specifically, for each section, we used other sections to train our model and predict gene expression for the section. All predictions were collected to assess the model performance.

### 2.2 The architecture of Hist2ST

This study proposed to predict the spatial gene expression from histology images through Hist2ST, a deep learning-based model. As shown in Fig 1, the input is the histology image that is divided into multiple patches around the sequenced spots. To process the input, Hist2ST consists of three modules: the Convmixer, Transformer, and graph neural network. The Convmixer module captures the 2D vision features within the image patch through convolution operations. The learned features are then fed into the Transformer module to obtain the spatial dependencies of all image patches (spots) through the self-attention mechanism. The learned features are input to the graph neural network module to explicitly learn the information from the nearest neighbours. Finally, the learned features are used to predict the gene expression by following the ZINB distribution.

#### 2.2.1 Convmixer module

The Convmixer module learns the 2D vision features within the image patch by convolution operations. Convmixer consists of a patch embedding layer followed by simple iterative applications of a convolutional block. Convmixer first extras the patch embeddings of each image patch by applying a convolution operation with *c_in_* input channels, *n* output channels, kernel size *p*, and stride *p*:

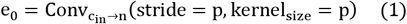

The core block of Convmixer consists of a depthwise convolution (i.e., grouped convolution whose group is equal to the number of channels) and a subsequent pointwise convolution (i.e., kernel size 1×1). Each convolution is followed by an activation function GLUE and post-activation BatchNorm as follows:

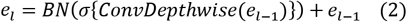

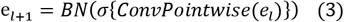

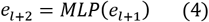

where *σ* and *BN* are the GLUE and BatchNorm, respectively. The MLP layer is used to project the learned 2D image patch into the one-dimensional feature vector of size 1024, which will be fed into the Transformer module.

#### 2.2.2 The Transformer module

Based on the learned features of spots from the Convmixer module, the Transformer module encodes the spatial locations of spots as follows:

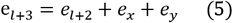

where e_*l*+2_ is outputted from the Convmixer module, and e_*x*_ ∈ *R*^1×1024^ and e_*y*_ ∈ *R*^1×1024^ are the encoded 2D coordinates of each spot through the function “torch.nn.Embedding” in Pytorch.

The multi-head attention layer in the Transformer module can be used to learn the attentions for a spots/image patch. The multi-head attention consists of multiple attention heads in a linear combination way as follows:

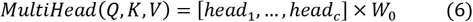

where *W*_0_ is the weight matrix for aggregating the attention heads, and *c* is the number of heads. Q, K, and V indicate Query, Key, and Value, respectively. The attention mechanism is defined as follows:

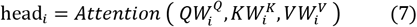

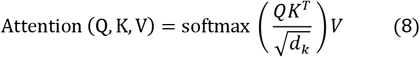

where 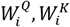, and 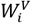 are weight matrixes. The term 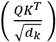 is called Attention Map, whose shape is N × N. The term V is the value of the self-attention mechanism, where *V* = *Q* = *K*. The attention weights contributed by other spots are stored in each column of the Attention Map. In the Transformer module, the dimension of the final output *h* is the same as the dimension of the input *e*_*l*+3_.

#### 2.2.3 The graph neural network module

The graph neural network is applied to explicitly learn the local spatial dependencies from the nearest neighboring spots. To be specific, we first construct the nearest neighboring graph, which then is fed into an inductive graph neural network GrahpSAGE [44] to learn the information from the neighboring spots.

##### Constructing the nearest neighboring graph

We use spatial locations in Euclidean distance as the intrinsic edges in the nearest neighboring graph due to the availability of spatial location in spots (image patch) [45]. Specifically, we select four nearest neighbors for each spot to construct the nearest neighboring graph *G* = (*V, E*). *N* = |*V*| means the number of spots, and *E* indicates the edges connecting with nearest neighbors. The Euclidean distance between every two spots *u* and *v* is calculated as follows:

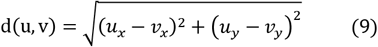

where *u_x_* and *u_y_* are the coordinates of node *u*.

##### The graph neural network module

The GraphSAGE framework is used to inductively learn information from the nearest neighbors. The input of GraphSAGE is the node feature matrix and the nearest neighboring graph. In our setting, the node is the spot with the corresponding feature *h* from the Transformer module, and the nearest neighboring graph is *G*. The GraphSAGE aggregates information of neighbors to generate node features for each spot. The aggregating layer of GraphSAGE can be formulated as follows:

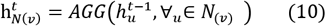

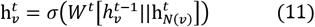

where N_(*v*)_ is the set of one-hop neighbors of spot *v*. The AGG is the mean () aggregator. Additionally, we follow the study [35] to use layer-aggregation mechanisms to avoid the over-smooth problem [46] through skip connections. We combine each layer of GraphSAGE by Long short-term memory (LSTM) [47] as follows:

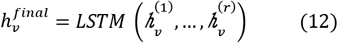

where 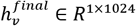 means the final features of spot *v*, and 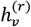 indicates the features of spot *v* in the *r* layer of the graph neural network. For all datasets, *r* = 4.

#### 2.2.4 The ZINB and MSE loss

After obtaining the features of each spot from the graph neural network module, we apply zero-inflated negative binomial (ZINB) layers to model the learned features. The ZINB model contains three separate full connection layers to estimate the parameters of ZINB: dropout rate π, dispersion degree θ, and mean μ.

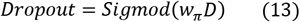

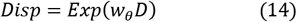

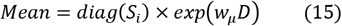

where *D* is the last layer of the graph neural network module. The S_*i*_ is the ratio of the total spot count to the median S. The ZINB model is parameterized by the ZINB distribution as follows:

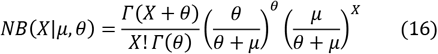

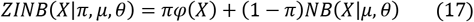

The loss function of ZINB is the sum of the negative log of ZINB likelihood of each data entry as follows:

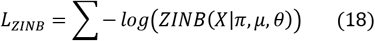

To learn the characteristic of gene expression, we also apply the Mean Square Errors loss as follows:

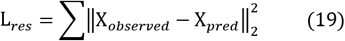

where *X_observed_* and *X_pred_* are the observed and predicted gene expression, respectively.

#### 2.2.5 Self-distillation mechanism

To alleviate the impact by the small spatial transcriptomics data, we used a similar self-distillation strategy to the previous study [48] to learn the “dark knowledge” from augmented samples. Specifically, for each image patch (anchor image patch), we generate five augmented image patches through random grayscale, rotation, and horizontal flip. Each anchor image patch and its augmented images are fed into our model and six predicted gene expressions will be outputted. We set learnable parameters for each predicted gene expression from the augmented image patch to learn their contribution to the final predicted gene expression as follows:

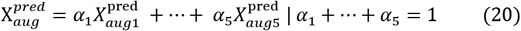

To fuse the knowledge information between the anchor image patch and the augmented image patches, we calculate the MSE loss as follows:

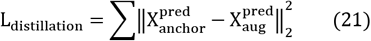

Finally, the total loss functions of our model are as follows:

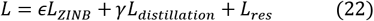

where *γ* and ϵ are the hyper-parameters to balance contribution from the self-distillation and ZINB. In this setting, we set *γ* = 0.5 and ϵ = 0.25 for all datasets.

### 2.3 Hyper-parameters setting

The Hist2ST was implemented in PyTorch and python. We set input channels *c_in_* =3, output channels n=32, kernel size p=7, and stride p=7 in the patch embedding layer of Convmixer. The kernel sizes of the depthwise and pointwise convolutional blocks were set to five and one, respectively. The input and output channels in depthwise and pointwise convolutional blocks were set to 32. For the Transformer module, the number of Multihead attention layers was eight, and the number of attention heads was sixteen. We set the number of layers of GNN as four and set the dimensions of both the input and hidden as1024. The number of nearest neighbors was set to four. The model was optimized through the Adam optimizer with a learning rate of 0.00001. We empirically set the number of epochs to 350 for all datasets. All results reported in this paper were conducted on Ubuntu 18.04.7 LTS with Intel® Core (TM) i7-8700K CPU @ 3.70 GHz and 256 GB memory.

For comparison, we ran the ST-Net and HisToGene methods with default parameters.

### 2.4 Evaluation criteria

#### 2.4.1 Pearson correlation coefficients

We measure the accuracies of predicted gene expression from histology images using the metric: Pearson correlation coefficients [49].

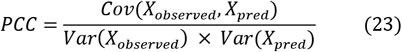

where Cov () is the covariance, and Var () is the variance. *X_observed_* ans *X_pred_* are the observed and predicted gene expression, respectively.

#### 2.4.2 Clustering performance

The commonly used metric Adjusted Rand Index (ARI) [50] is applied for evaluating clustering results in this study.

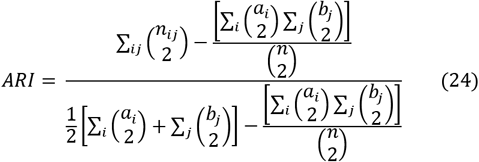

where a_*i*_ and b_*j*_ are the number of cells appearing in the i-th cluster of predicted clusters C and the j-th cluster of true clusters Y, respectively. n_*ij*_ means the number of overlaps between the i-th cluster of C and the j-th cluster of Y.

## 3. Results

### 3.1 Performance on the spatial transcriptomics datasets

We assessed all models on the HER2+ (containing 32 tissue sections) and cSCC (containing 12 tissue sections) spatial transcriptomics datasets through the leave-one-out cross validation. As shown in Fig 2, our method achieved superior results in terms of the mean Pearson correlation coefficients between the observed and predicted genes. To avoid outliers, we also showed the median Pearson correlation coefficients and that our method consistently outperformed other methods. Concretely, the mean Pearson correlation coefficients of our method were 9% and 11% higher than the second-ranked method HisToGene on HER2+ and cSCC datasets, respectively. Each tissue section of our method had a higher Pearson correlation coefficient than the competing methods. Relatively, our method achieved the best results on B1-6 tissue sections among all tissue sections of the HER2+ dataset. All methods performed low on E1-3 and F1-3 tissue sections, whereas our method still performed better than the second-ranked method HisToGene (HisToGene was 8% smaller than our method in terms of mean Pearson correlation coefficients on these tissue sections). ST-Net had the lowest performance on most tissue sections in the HER2+ and cSCC datasets. These results indicated that our model could efficiently predict the gene expression pattern from the histology images.

**Fig 2.**
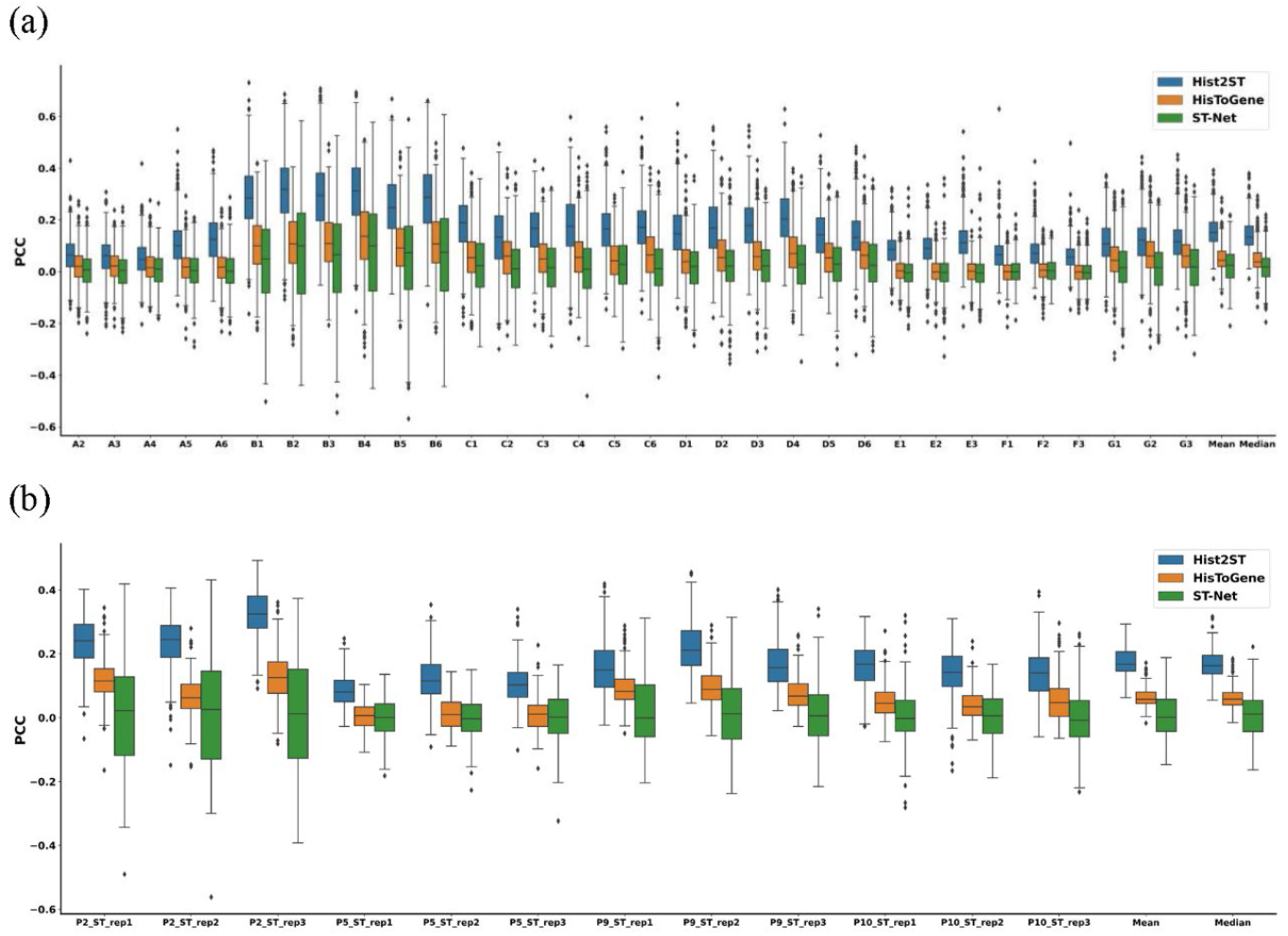
Evaluation of gene expression prediction on the (a) HER2+ and (b) cSCC datasets by the Pearson correlation coefficients between the observed and predicted gene expressions by three methods.

To investigate the contributions of components for our method Hist2ST, we conducted ablation studies on spatial transcriptomics datasets. As shown in Fig 3, the removal of any module in our model caused performance decreases. Specifically, the removal of the Convmixer module caused a decrease of 4% in terms of average Pearson correlation coefficients. This change indicated that the 2D vision features in the image patch captured by Convmixer could help the model learn the pattern of gene expression. The results of our method without the module Convmixer were better than those of HisToGene due to the inclusion of the graph neural network. The removal of the Transformer module caused a decrease of 11% in terms of average Pearson correlation coefficients. The results of our method without the module Transformer were similar to those of ST-Net. The results indicated that the Transformer module efficiently captured the global relationships between image patches. The removal of the graph neural network module caused a small but significant drop (2% in terms of average Pearson correlation coefficients), indicating the useful information learned from nearest neighbors helped our method to capture the gene expression from histology images. The removal of the self-distillation mechanism caused a decrease of 1% in terms of the average Pearson correlation coefficients, which indicated that the self-distillation mechanism could relieve the overfitting problem due to limited training datasets. The trend was similar for the cSCC dataset (Fig S1). In summary, the better performance of our model relied on the cooperation of the modules.

**Fig 3.**
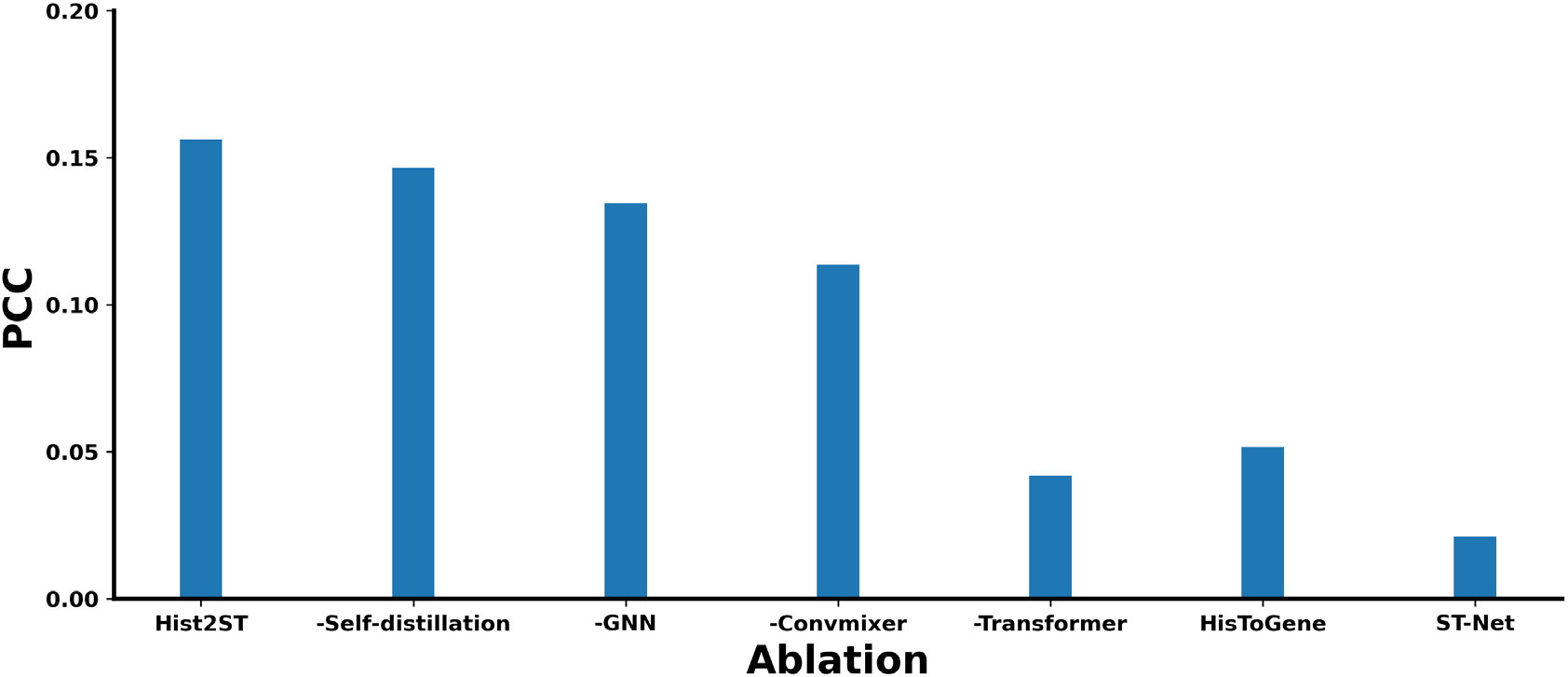
The results on the HER2+ dataset of our method by excluding an individual module.

### 3.2 Visualization of the predicted gene expression

To further understand the predicted gene expression, we visualized the top predicted genes on the histology image. Each predicted gene was calculated by the Pearson correlation coefficients and was ranked by the average -log10 p-value across all tissue sections (Table S1). For the HER2+ dataset, we visualized the top four genes (FN1, GANS, SCD, and MYL12B) on the tissue sections with the smallest p-value. As shown in Fig 4, the visualized results showed that the predicted genes of our model achieved the highest Pearson correlation coefficients on these top four genes. Compared to other methods, our method had more similar gene expression patterns to the observed gene expression. Additionally, all these four top predicted genes were the maker genes in breast cancer [29, 51–53]. For example, the previous study [53] had shown that the high expression levels of COL1A1 and FN1 correlated to an advanced stage of breast cancer and poor clinical outcomes. As shown in Fig S2 and Table S2, we also took a similar strategy to visualize the four top genes (MSMO1, NDRG1, ITGA6, and DMKN) on the tissue sections with the smallest p-values on the cSCC dataset. We found these four top genes were the maker genes reported in the literature [54–56]. Our method consistently outperformed competing methods in terms of average Pearson correlation coefficients on these genes. The results indicated that our model could accurately predict the gene expression and reserve maker genes information. As a comparison, we also showed the top four genes predicted by the second-ranked method HisToGene on HER+2 and cSCC datasets. As shown in Fig S3 and Table S3, the mean Pearson correlation coefficients of the top four genes (GANS, FASN, MYL12B, and SCD) predicted by HisToGene were much lower than the top four genes predicted by our method on the HER+2 dataset. Meanwhile, our method still outperformed HisToGene in terms of average Pearson correlation coefficients of these four top genes predicted by HisToGene. The trend was similar for the cSCC dataset (Fig S4 and Table S4).

**Fig 4.**
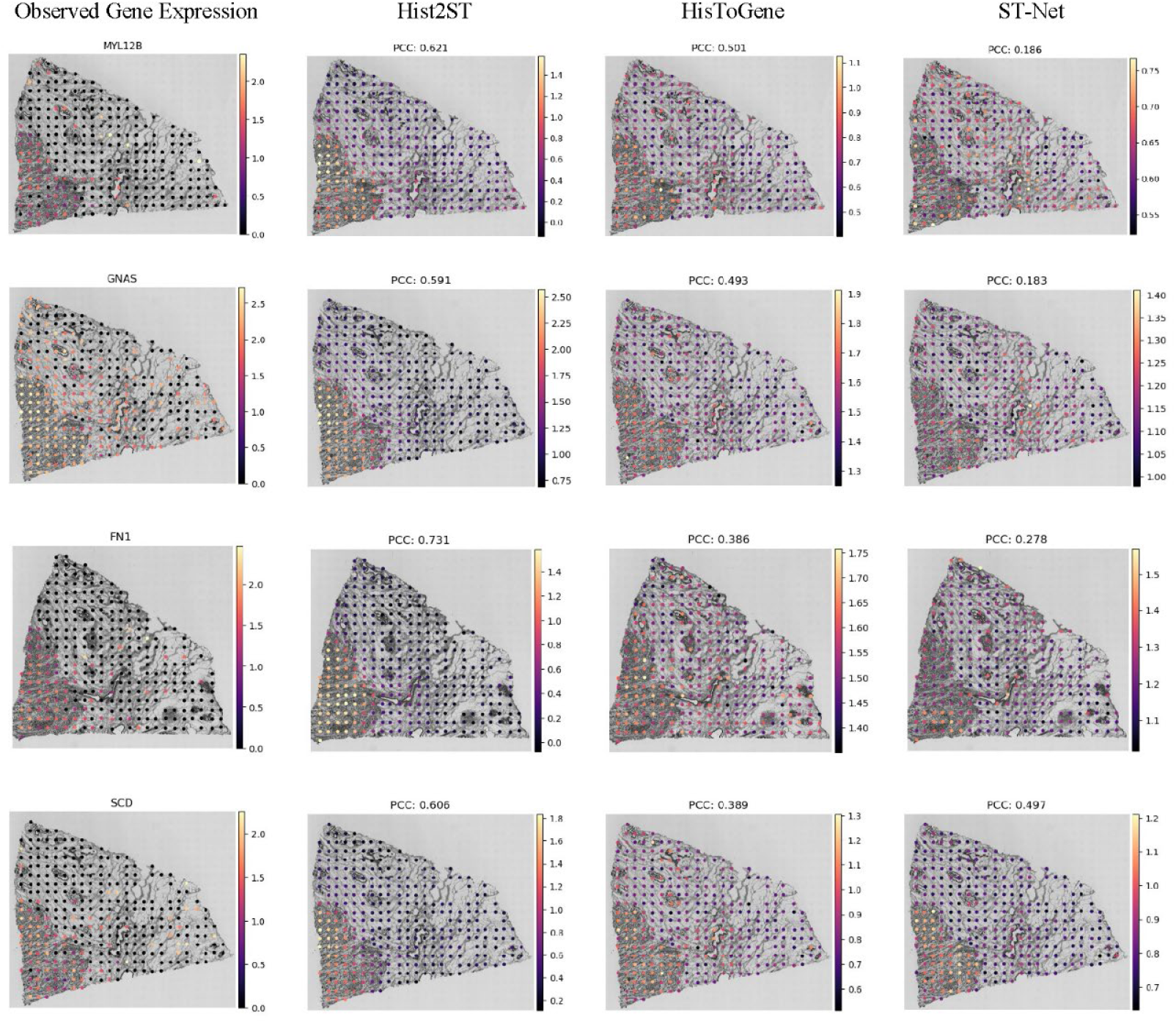
Visualization of the HER2+ dataset by the top four predicted genes with the highest values of average -log10 p-values across all tissue sections, where the p-value for each tissue section was obtained according to the correlation between the predicted and observed gene expression. For each of the four genes, the tissue section that had the smallest p-value by our model was selected for visualization.

To show how well our method can retain the biological signals, we used the R package clusterprofile [57] to conduct gene set enrichment. Inspired by IPath [58], we ranked the genes by the average -log10 p-values across all tissue sections, and then selected the top 100 genes to conduct the gene enrichment by clusterprofile. For the HER2+ dataset, the highly correlated predicted genes were shown to be enriched in breast cancer-related pathways (Fig S5). For instance, the top enriched pathway, the NADH dehydrogenase activity, was reported to profoundly enhance the aggressiveness of human breast cancer cells in the recent literature [59].

### 3.3 Spatial region detection

To evaluate the performance of each method in detecting spatial regions in the whole histology image, we performed K-means clustering using the predicted gene expression from each method. Since only the HER2+ dataset had six tissue sections (B1, C1, D1, E1, F1, and G2) with pathologists’ annotation, we compared each method using these tissue sections. As a comparison, we also used the observed gene expression to conduct clustering analyses. The clustering results were evaluated through ARI by regarding the pathologist’s annotated spatial regions as the ground truth. We expected the identified spatial clusters using the predicted gene expression to accord with the pathologist’s annotated spatial regions. As shown in Fig 5, our model achieved the highest average ARI on six tissue sections. Concretely, the average ARI of our model was 7% higher than the second-ranked method HisToGene. Our method obtained the highest ARIs for four sections (B1, C1, D1, and F1). Both our method and HisToGene performed low in terms of the E1 tissue section. We found that the clustering results achieved using the observed gene expression of tissue section E1 were similar to those of our method. It is likely that the sequenced gene expressions contained noise raised from sequencing techniques. Our model also was 5% higher than the observed gene expression in terms of average ARI. It is likely that the gene expression learned by our model contained the extra image information. HisToGene achieved a similar average ARI to the observed gene expression. ST-Net achieved the highest ARI on tissue section E1 while it performed low on other tissue sections. Relatively, the spatial regions identified using gene expression predicted by our method were more consistent with those annotated by pathologists than other methods.

**Fig 5.**
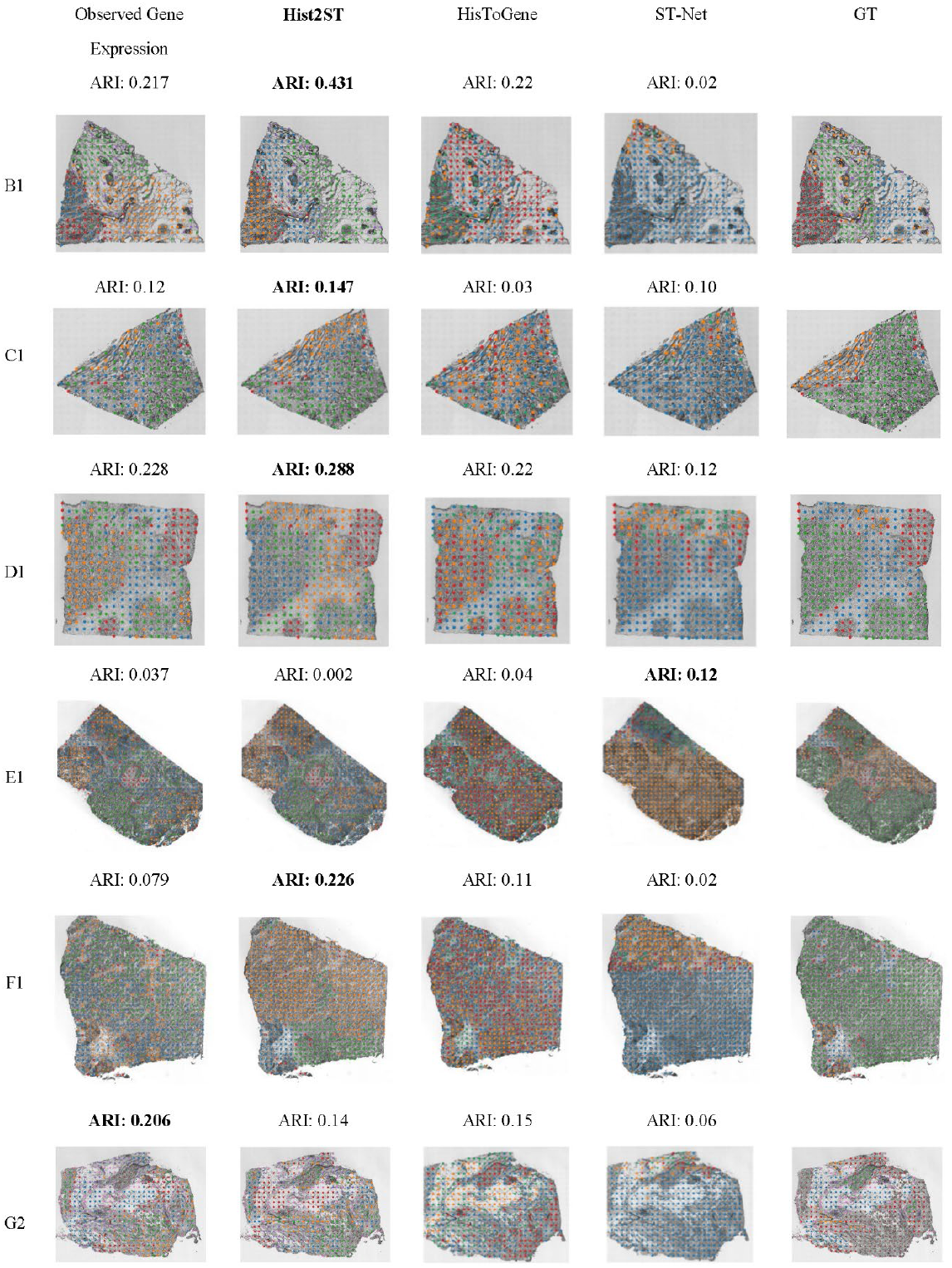
Spatial region detection on the HER2+ dataset using gene expressions obtained by all methods, where the observed represents the direct use of sequenced gene expressions, and the GT represents the ground truth labels from the pathology annotations. The results of HisToGene and ST-Net were directly obtained from the reference [27].

## 4. Discussion

The spatial transcriptomics technologies hold the potential of profiling gene expression for the whole transcriptomics at nearly single cell resolution accompanied with their spatial locations, for many techniques, and often with the matched whole slide of hematoxylin and eosin (H&E)- stained histology images (WSI). Relatively, WSIs are cheap and easy to obtain and are routinely generated in clinics. It is desirable to predict the gene expression from WSI. Here, we have developed Hist2ST, a deep learning model that uses histology images to predict spatial gene expression. At each sequenced spot, the corresponding histology image is cropped into an image patch, from which 2D vision features are learned by the module Convmixer through convolutional operations. Meanwhile, the spatial relations with the whole image and neighbored patches are captured through Transformer and graph neural network modules, respectively. The learned features are then used to predict the gene expression by following the ZINB distribution. Hist2ST was tested on the HER2+ breast cancer and the cutaneous squamous cell carcinoma datasets, and shown to outperform existing methods in terms of both gene expression prediction and spatial region identification. Thus, Hist2ST enables generating spatial transcriptomics data from histology images for elucidating molecular signatures of tissues.

Compared to ST-Net and HisToGene, Hist2ST benefits from the consideration of both the 2D vision features and the spatial dependency. HisToGene includes the spatial locations through Transformer while ignoring the 2D vision features within the image patch. However, our model can learn the 2D vision features by the Convmixer module through convolution operations. ST-Net captures the 2D vision feature while ignoring spatial locations. Our model considers spatial locations by using the Transformer and graph neural network modules. On the other hand, to alleviate the impact by the small spatial transcriptomics data, a self-distillation mechanism is employed for the efficient learning of our model. Benefiting the above dedicated design, our model achieves the best performance compared to the state-of-the-art methods. Additionally, our method achieves similar inference times (about second seconds for each tissue section) to competing methods. The pathway analyses indicate that the gene expression predicted by our model also reserves biological meaningful information.

Despite the advantages of Hist2ST, our model can be improved in several aspects. For example, the better performance of the deep learning based-model would need a relatively large training set, such as Hist2ST. We adapt the data efficiency learning way by self-distillation technology to alleviate the limited training datasets. On the other hand, with the development of ST, more and more training ST data can be available in the near future. In summary, we demonstrate that Hist2ST provides a deep learning model for generating spatial transcriptomics data from histology images for elucidating molecular signatures of tissues.

## Key Points

- The histology images are relatively easy and cheap to obtain, so it is promising to predict spatial gene expression from histology images. Current methods for predicting gene expression from histology depended on either the 2D vision features or the spatial dependency, but didn’t combine these together.
- We have developed Hist2ST, a deep learning-based model that predicts spatial RNA-seq expression from histology images. Hist2ST simultaneously considers the 2D vision features and the spatial dependency through convolutional operations, Transformer, and graph neural network modules. The learned features are used to predict the gene expression by following the ZINB distribution.
- Hist2ST was tested on the HER2+ breast cancer and cutaneous squamous cell carcinoma datasets, and shown to outperform existing methods in terms of both spatial gene expression prediction and spatial region identification.

## Code availability

All source codes used in our experiments have been deposited at https://github.com/biomed-AI/Hist2ST

## Data availability

The spatial transcriptomics datasets that support the findings of this study are available here: (1) human HER2-positive breast tumor ST data https://github.com/almaan/her2st/. (2) human cutaneous squamous cell carcinoma 10x Visium data (GSE144240).

## Funding

This study has been supported by the National Key R&D Program of China (2020YFB0204803), National Natural Science Foundation of China (61772566), Guangdong Key Field R&D Plan (2019B020228001 and 2018B010109006), Introducing Innovative and Entrepreneurial Teams (2016ZT06D211), Guangzhou S&T Research Plan (202007030010).

## Notes

### Competing Interest Statement

The authors have declared no competing interest.

